# Pervasive environmental chemicals impair oligodendrocyte development

**DOI:** 10.1101/2023.02.10.528042

**Authors:** Erin F. Cohn, Benjamin L.L. Clayton, Mayur Madhavan, Sara Yacoub, Yuriy Federov, Katie Paul-Friedman, Timothy J. Shafer, Paul J. Tesar

**Affiliations:** Department of Genetics and Genome Sciences, Case Western Reserve University School of Medicine, Cleveland, Ohio 44106, USA; Center for Computational Toxicology and Exposure, Office of Research and Development, U.S. Environmental Protection Agency, Research Triangle Park, North Carolina 27711, USA

## Abstract

Exposure to environmental chemicals can impair neurodevelopment^1–4^. Oligodendrocytes that wrap around axons to boost neurotransmission may be particularly vulnerable to chemical toxicity as they develop throughout fetal development and into adulthood^5,6^. However, few environmental chemicals have been assessed for potential risks to oligodendrocyte development. Here, we utilized a high-throughput developmental screen and human cortical brain organoids, which revealed environmental chemicals in two classes that disrupt oligodendrocyte development through distinct mechanisms. Quaternary compounds, ubiquitous in disinfecting agents, hair conditioners, and fabric softeners, were potently and selectively cytotoxic to developing oligodendrocytes through activation of the integrated stress response. Organophosphate flame retardants, commonly found in household items such as furniture and electronics, were non-cytotoxic but prematurely arrested oligodendrocyte maturation. Chemicals from each class impaired human oligodendrocyte development in a 3D organoid model of prenatal cortical development. In analysis of epidemiological data from the CDC’s National Health and Nutrition Examination Survey, adverse neurodevelopmental outcomes were associated with childhood exposure to the top organophosphate flame retardant identified by our oligodendrocyte toxicity platform. Collectively, our work identifies toxicological vulnerabilities specific to oligodendrocyte development and highlights common household chemicals with high exposure risk to children that warrant deeper scrutiny for their impact on human health.

## MAIN

Humans are exposed to a plethora of environmental chemicals with unknown toxicity profiles. The developing central nervous system is particularly sensitive to environmental insults and chemical exposures can be especially harmful to children if they occur during critical periods of development^1,2^. For example, the heavy metals methylmercury and lead, as well as industrial chemicals such as polychlorinated biphenyls are known to disrupt brain development^3,4^.Chemical exposures may trigger pathogenesis or exacerbate underlying genetic factors^7–9^. Importantly, the prevalence of neurodevelopmental disorders including autism spectrum disorder and attention-deficit hyperactivity has increased^10,11^, however, genetic factors can account for less than half of cases^2^. Therefore, evaluating how environmental factors, including chemical exposures, contribute to or initiate neurodevelopmental disorders has become imperative.

While neurons have been more thoroughly evaluated for their susceptibility to chemical toxicity^12,13^, non-neuronal or glial cells are also essential for normal brain function. Oligodendrocytes generate myelin, a requirement for efficient neuronal transmission, and provide metabolic trophic support to neurons which is essential for their function and longevity^5,14^. Conversely, impaired oligodendrocyte development or their loss results in significant cognitive and motor disability in genetic diseases such as Pelizaeus-Merzbacher disease, and in inflammatory diseases such as multiple sclerosis^15–17^. A few environmental chemicals including natural products and industrial compounds reportedly alter oligodendrocyte function^18,19^. However, the vast majority of chemicals present in the environment have not been evaluated for oligodendrocyte toxicity, in large part, due to prior challenges in capturing oligodendrocyte development at high purity and scale.

Both oligodendrogenesis and myelination have wide windows of vulnerability for environmental chemical exposure in humans. Development of oligodendrocytes from oligodendrocyte progenitor cells (OPCs) occurs throughout the first two years of life, and myelination begins during fetal development, peaks in infancy and childhood, and continues into adolescence and adulthood^6,20^. Therefore, oligodendrocytes are not only vulnerable during fetal development but also long after birth. In this study, we developed a toxicity screening platform to assess 1,823 chemicals that belong to the rapidly expanding repertoire of environmental contaminants. We identified chemicals belonging to two classes commonly found in households that perturb oligodendrocyte development.

## RESULTS

### Phenotypic screen for environmental chemicals that disrupt oligodendrocyte development

Previously, we established methods to generate OPCs from mouse pluripotent stem cells (mPSCs) at the scale required for high-throughput screening efforts^21–23^. mPSC-derived OPCs reliably develop into oligodendrocytes over 3 days *in vitro*, providing a robust approach for identifying environmental chemicals that affect oligodendrogenesis. We screened a library of 1,823 chemicals to assess their effects on development of OPCs into oligodendrocytes (Fig. 1a). This library contains diverse chemicals with the potential for human exposure, including industrial chemicals, pesticides, and chemicals that are of interest to regulatory agencies^24^. In a primary screen, we treated OPCs with chemicals at a concentration of 20 μM and allowed oligodendrocytes to develop for 3 days before analysis. Toxicity screening during the OPC to oligodendrocyte developmental transition enabled us to identify both chemicals cytotoxic to developing oligodendrocytes and chemicals that impede oligodendrocyte generation without being cytotoxic.

**Fig. 1:**
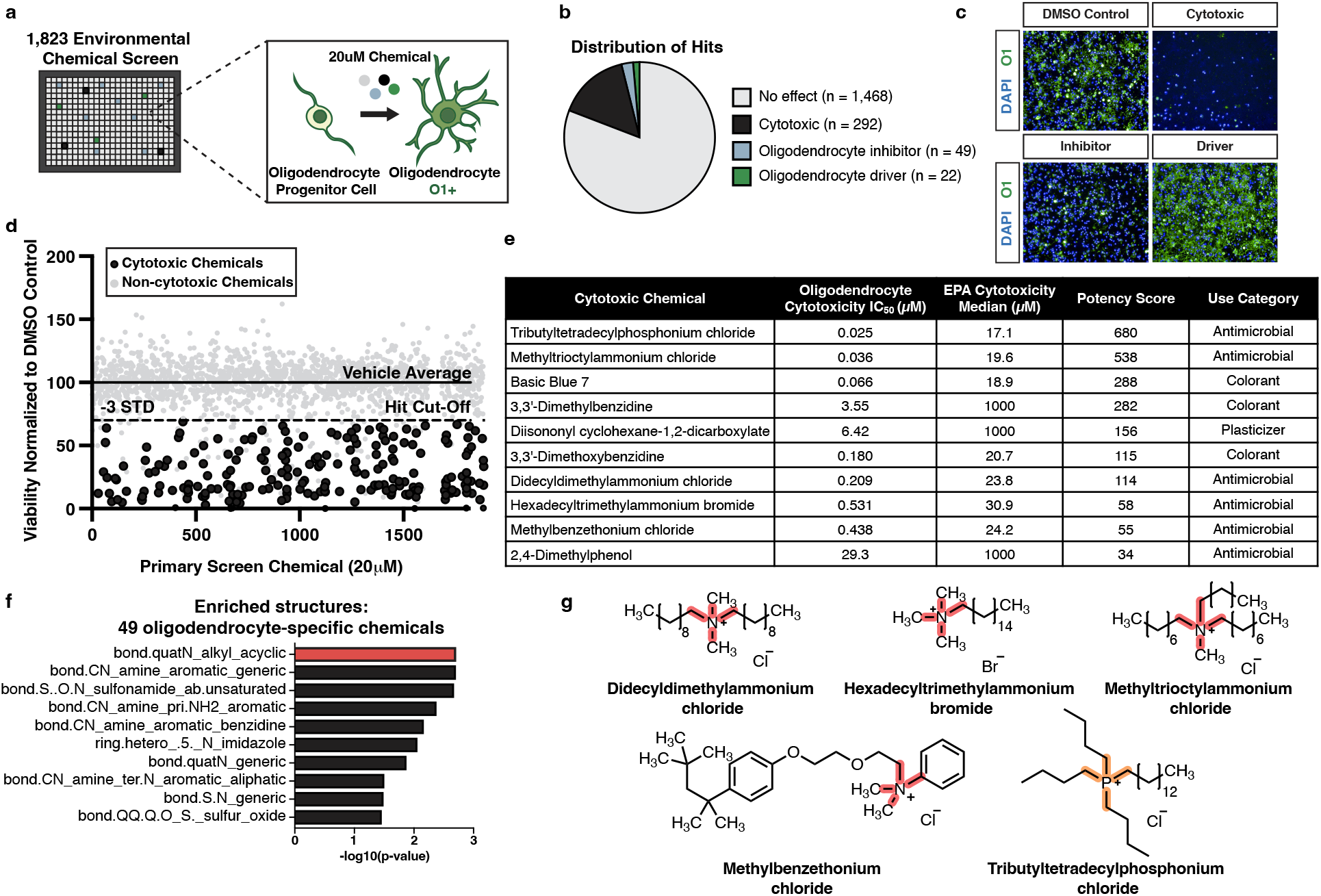
Quaternary compounds are potently cytotoxic to developing oligodendrocytes. **a,** Schematic of the primary chemical screen in mPSC-derived oligodendrocytes. **b,** Pie chart of the number of cytotoxic chemicals (black), inhibitors of oligodendrocyte development (blue), and drivers of oligodendrocyte development (green) identified from the primary chemical screen, along with chemicals that had no effect (gray). **c,** Representative immunohistochemistry images after 3 days of oligodendrocyte development. Each image shows cells cultured with DMSO (vehicle control), or one of three chemicals with different effects on oligodendrocyte generation. Nuclei are marked using DAPI (in blue) and oligodendrocytes are marked using O1 (in green). **d,** Primary chemical screen showing the effect of 1,823 environmental chemicals on the viability of developing oligodendrocytes displayed as viability normalized to vehicle control. The solid line represents the average of the vehicle control set at 100%. The dotted line marks a reduction in viability of 30% (>3 standard deviations). The 206 cytotoxic hits that pass this threshold and were validated by MTS are colored in black. Non-cytotoxic chemicals and cytotoxic hits not validated by MTS are colored in gray. **e,** Table showing characteristics of 49 oligodendrocyte-specific cytotoxic hits tested in 10-point dose response from (40 nM to 20 μM). IC50 values were determined with curvefitting and compared to median cytotoxicity values obtained from the EPA database for each chemical. Potency scores were calculated by dividing the cytotoxicity median by the experimentally determined IC50 in oligodendrocytes. Chemicals were ranked based on increasing potency score. Table also includes each chemical’s use category. **f,** Chemotype analysis for the 49 oligodendrocyte-specific cytotoxic compounds, with the most enriched structural domain, bond.quatN_alkyl_acyclic (p-value = 0.002, OR = 16.2), highlighted in red. p-values were generated using a one-sided Fisher’s exact test. **g,** Chemical structures for the four quaternary ammonium compounds and one quaternary phosphonium compound. The enriched cytotoxicity-associated bond for quaternary ammonium compounds is highlighted in red. The quaternary phosphonium bond is highlighted in orange.

We determined chemical cytotoxicity in oligodendrocytes by quantifying viable nuclei based on staining with 4’,6-diamidino-2-phenylindole (DAPI) and considered chemicals cytotoxic that reduced viability by more than 30% compared to the negative control. We further classified the remaining non-cytotoxic chemicals based on whether they interfered with the development of oligodendrocytes from OPCs by immunostaining for the O1 antigen, which is exclusively expressed on maturing oligodendrocytes. We considered non-cytotoxic chemicals that reduced the percentage of O1-positive cells by greater than 50% to be inhibitors of oligodendrocyte development. Conversely, we considered chemicals that increased the percentage of O1-positive cells by more than 20% to be drivers of oligodendrocyte development (Fig. 1b,c, and Extended Data Fig. 1a,b). Of the 1,823 chemicals in the primary screen, more than 80% had no effect on oligodendrocyte development or cytotoxicity, 292 were identified as cytotoxic to developing oligodendrocytes, 49 inhibited oligodendrocyte generation, and 22 stimulated oligodendrocyte generation (Fig. 1b, and Supplementary Table 1).

### Quaternary compounds are selectively and potently cytotoxic to oligodendrocyte development

To validate cytotoxic hits from the primary screen we used a colorimetric MTS tetrazolium assay designed to assess cell viability by measuring metabolic activity (Fig. 1d). To identify chemicals with specific cytotoxicity to oligodendrocyte development, we compared cytotoxicity profiles of 206 MTS-validated chemicals from the primary screen to both an in-house primary screen in mouse astrocytes, representing another glial subtype, and a public database from the US Environmental Protection Agency (EPA), that contains cytotoxicity data for many cell types but not glial cells (Fig. 1e and Extended Data Fig. 2b). Chemicals cytotoxic to oligodendrocyte development but non-cytotoxic to astrocytes were tested in 10-point dose response (40 nM to 20 μM), and used to calculate IC50 values for each chemical (Fig. 1e and Extended Data Fig. 2c). Finally, we ranked the top ten cytotoxic chemicals based on potency in developing oligodendrocytes, lack of cytotoxicity to astrocytes, and lack of potency in cytotoxicity assays using other cell types (Fig. 1e).

Through computational analysis we identified a chemical structure, characterized by a central nitrogen with four alkyl groups (bond.quatN_alkyl_acylic), as the most enriched structural domain among chemicals cytotoxic specifically to oligodendrocytes. This bond defines 13 quaternary ammonium compounds in the 1,823 chemical library. The primary screen identified 9 of these chemicals as cytotoxic to oligodendrocyte development, four of which are found in the top 12 cytotoxic hits, including methyltrioctylammonium chloride (Fig. 1f,g, and Supplementary Table 2). The most highly ranked oligodendrocyte cytotoxic chemical, tributyltetradecylphosphonium, a quaternary phosphonium compound, has similar structure and function to quaternary ammonium compounds^25^. Given that our primary screen and secondary validation assays utilized mPSC-derived oligodendrocytes, we next confirmed cytotoxicity for two of the top ranked cytotoxic hits on oligodendrocytes generated from primary OPCs isolated directly from mouse postnatal brain tissue. Both methyltrioctylammonium chloride and tributyltetradecylphosphonium chloride, the two most highly ranked quaternary ammonium and phosphonium compounds (quaternary compounds) were cytotoxic at 20 μM to primary OPCs, resulting in greater than 80% reduction in cell viability (Extended Data Fig. 2d,e). Collectively, these data demonstrate that quaternary compounds are cytotoxic to developing oligodendrocytes. While quaternary phosphonium compounds are an emerging class of disinfectants^25^, current human exposure to quaternary ammonium compounds is likely as they are widely used in cosmetic products and as disinfecting agents, an application that increased significantly due to the COVID-19 pandemic^26^.

### Quaternary compounds activate the integrated stress response to induce apoptosis

To identify the mechanism underlying quaternary compound cytotoxicity in developing oligodendrocytes, we performed RNA sequencing on cells treated for 4 hours with two quaternary compounds: tributyltetradecylphosphonium chloride and methyltrioctylammonium chloride. Gene set enrichment analysis (GSEA) revealed that quaternary compound exposure results in enrichment for hallmark gene sets involved in programmed cell death and the integrated stress response (ISR) (Fig. 2a,b, and Supplementary Table 3). The ISR is activated by diverse environmental stressors and if not resolved, can lead to cell death. We confirmed activation of the ISR by performing qPCR using C/EBP homologous protein (CHOP), as a canonical marker of ISR activation and mediator of ISR-induced apoptosis (Fig. 2c). To determine the mechanism of cell death following ISR activation, we screened small molecule inhibitors of multiple programmed cell death pathways. We found that only QVD-OPH, an inhibitor of apoptosis, was able to prevent cell death induced by exposure to quaternary compounds (Extended Data Fig. 2f). These data suggest that quaternary compounds initiate ISR-mediated apoptosis in developing oligodendrocytes.

**Fig. 2:**
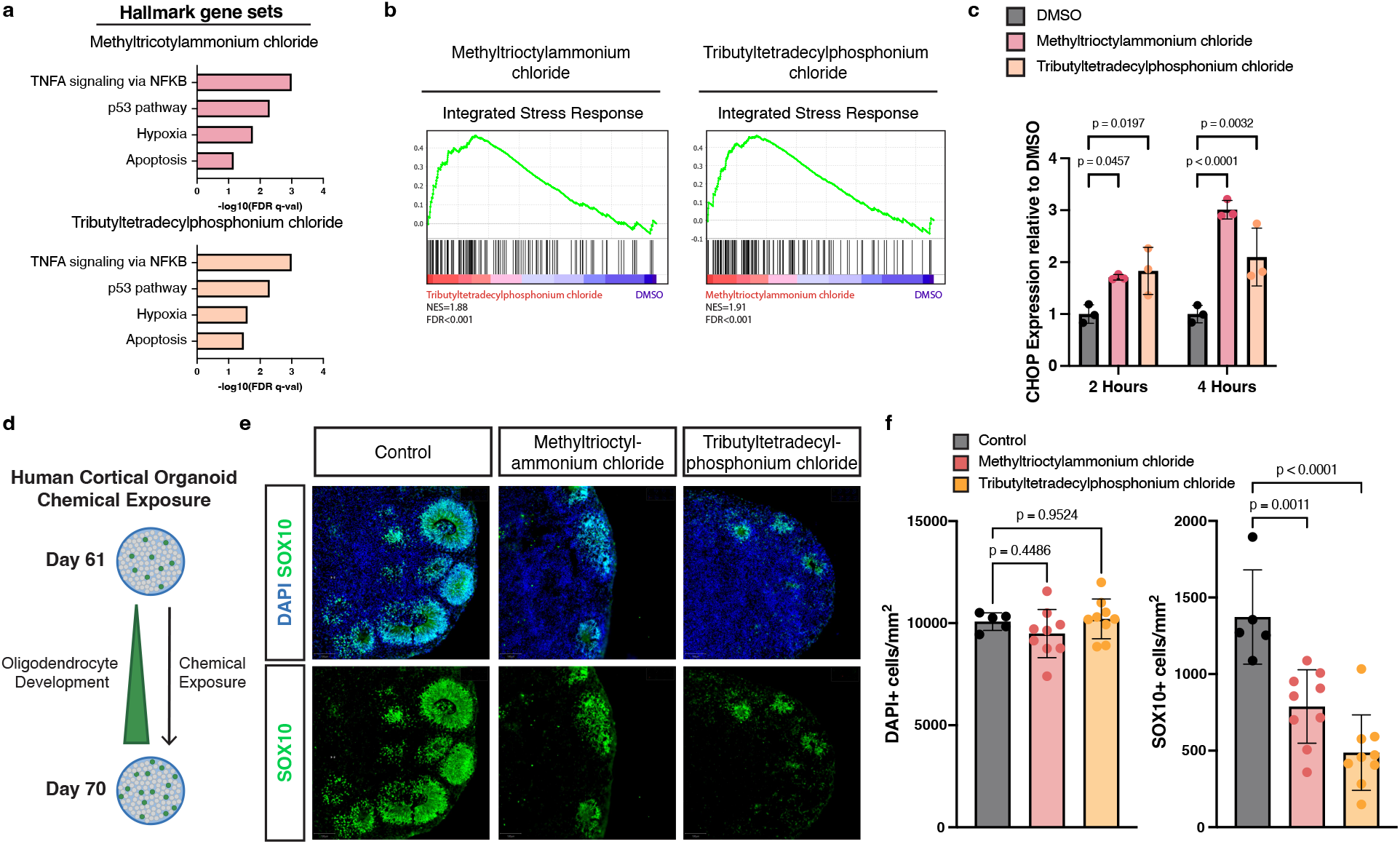
Quaternary compounds activate the integrated stress response and are cytotoxic to human oligodendrocytes. **a,** Gene set enrichment analysis (GSEA) of hallmark gene sets upregulated in OPCs in response to incubation with 20 μM quaternary ammonium (red) or phosphonium compounds (orange) for 4 hours. **b,** GSEA of an integrated stress response gene set in OPCs treated with methyltrioctylammonium chloride or tributyltetradecylphosphonium chloride compared with DMSO treated OPCs demonstrates significant enrichment (FDR<0.001) for genes involved in the integrated stress response (normalized enrichment scores [NES] = 1.91, 1.88 respectively). **c,** qRT-PCR of CHOP in OPCs treated with DMSO (gray), methyltrioctylammonium chloride (red), and tributyltetradecylphosphonium chloride (orange). Data are presented as the mean value ± standard deviation from three biological replicates, represented as closed circles. P values were calculated using one-way ANOVA with Dunnett post-test correction for multiple comparisons. **d,** Schematic depicting exposure of human cortical organoids to cytotoxic chemicals. **e,** Representative immunohistochemistry images of control human cortical organoids and organoids treated for 10 days with 360 nM methyltrioctylammonium chloride or 300 nM tributyltetradecylphosphonium chloride (approximate IC_90_ in mPSC-derived oligodendrocytes). Images show all cells (DAPI+, blue) and oligodendrocytes (SOX10+, in green) at day 70. **f,** Quantification of total cell number (DAPI+ per mm2) and oligodendrocytes (SOX10+ per mm2) in the whole cortical organoid. Data are presented as the mean value ± standard deviation from n ≥ 5 biological replicates (individual organoids) indicated by closed circle data points. p-values were calculated using one-way ANOVA with Dunnett post-test correction for multiple comparisons.

### Quaternary compounds are cytotoxic to human oligodendrocyte development in cortical organoids

To determine whether quaternary compounds could disrupt human oligodendrocyte development, we leveraged our human pluripotent stem cell (hPSC)-derived regionalized neural organoid model in which oligodendrogenesis and myelination are integrated with fundamental processes of prenatal cortical development^27,28^. We supplemented media with methyltrioctylammonium chloride and tributyltetradecylphosphonium chloride on day 60, a critical time point for oligodendrocyte development (Fig. 2d). After culturing organoids in the presence of quaternary compounds for 10 days, we harvested organoids for analysis. Given that cell density was maintained across all conditions we concluded that quaternary compounds are not broadly cytotoxic. However, in immunohistochemistry analysis using the oligodendrocyte lineage marker, SOX10, we documented a significant reduction in SOX10-positive OPCs and oligodendrocytes (Fig. 2e,f). These results suggest that quaternary compounds are cytotoxic to developing human oligodendrocytes in an *in vitro* model of early human brain development.

### Organophosphate flame retardants arrest oligodendrocyte development

Many toxicity screens evaluate cell viability as a single endpoint measure. However, our screening platform allowed us to both identify cytotoxic chemicals and evaluate whether non-cytotoxic chemicals affect an essential developmental transition. Of the 1,539 non-cytotoxic compounds identified in the primary screen, 71 altered oligodendrocyte development. The 22 chemicals identified as enhancers of oligodendrocyte development largely consisted of thyroid hormone receptor modulators which are well known to drive oligodendrocyte generation (Extended Data Fig. 3a)^29^. The remaining 49 chemicals inhibited oligodendrocyte development (Fig. 3a). Computational analysis of oligodendrocyte inhibitors revealed an enriched structure characterized by a central phosphate (bond.P.O_phosphate_alkyl_ester) as a top enriched structure with the highest odds ratio (Fig. 3b, and Supplementary Table 2). This structure is found in three chemical hits, tris(methylphenyl) phosphate (TMPP), tris(2,3-dibromopropyl) phosphate (TBPP), and tris(1,3-dichloro-2-propyl) phosphate (TDCIPP). These chemicals are all organophosphate esters that belong to a large class of compounds widely used as both pesticides and flame retardants (Fig. 3c, and Extended Data Fig. 3b). Of the 13 organophosphate flame retardants in the primary screen chemical library, 7 chemicals reduced the percentage of O1-positive oligodendrocytes. We tested the top 3 organophosphate flame retardants in 8-point dose response (30 nM to 20 μM) and used these data to generate IC50 values (Fig. 3d, and Extended Data Fig. 3b). All three organophosphate flame retardants also inhibited the development of oligodendrocytes from mouse OPCs isolated directly from postnatal brain tissue (Extended Data Fig. 3c,d).

**Fig. 3:**
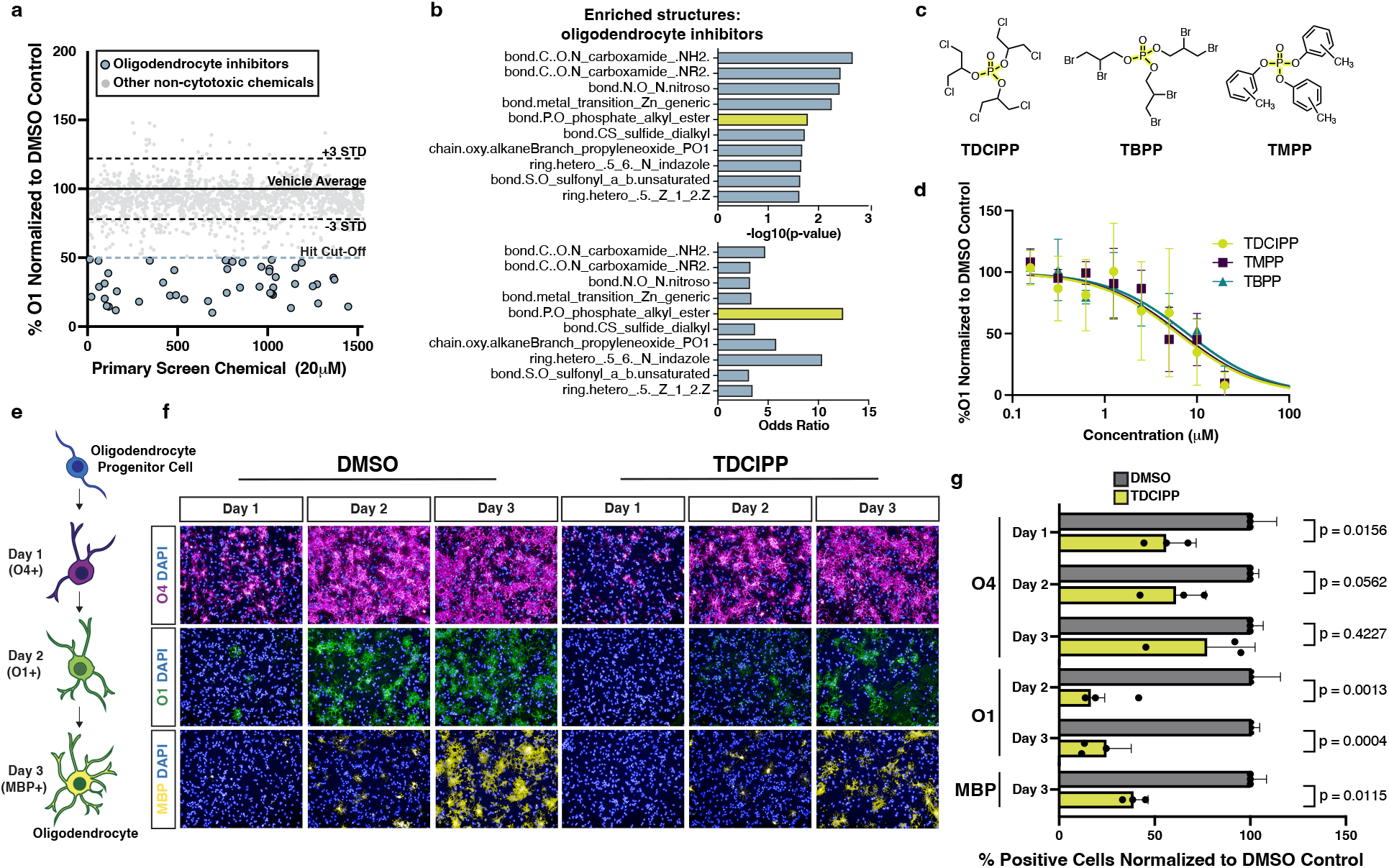
Organophosphate flame retardants arrest oligodendrocyte maturation. **a,** Primary chemical screen showing the effect of 1,539 non-cytotoxic environmental chemicals on oligodendrocyte development displayed as percent O1+ cells normalized to DMSO. The dotted lines mark ± 3 standard deviations from the mean of control wells. The blue dotted line marks the inhibitor hit cutoff, a reduction in O1+ cells of 50% (>7 standard deviations) compared to DMSO. The 49 oligodendrocyte inhibitors that pass this threshold are colored in blue. All other non-cytotoxic chemicals that did not inhibit oligodendrocyte development are colored in gray. **b,** Chemotype analysis for oligodendrocyte inhibitors showing both the p-value and odds ratio. Among the top most significant structural domains, bond.P.O_phosphate_alkyl_ester (p-value = 0.02, OR = 12.5) has the highest odds ratio, and is highlighted in yellow. p-values were generated using a one-sided Fisher’s exact test. **c,** Chemical structures for three organophosphate flame retardants containing the structure bond.P.O_phosphate_alkyl_ester, highlighted in yellow. **d,** Graph of eight-point dose response (30 nM to 20 μM) quantifying the effect of three organophosphate flame retardants on oligodendrocyte (O1+) generation from OPCs. Data are presented as the mean value ± standard deviation from three biological replicates (OPC batches generated from independent mPSC lines). **e,** Schematic showing stages of in vitro oligodendrocyte development and the markers for early (O4), intermediate (O1), and late (MBP) oligodendrocytes. **f,** Representative images of early (O4+, in magenta), intermediate (O1+, in green), and late (MBP+ in yellow) oligodendrocytes after treatment with DMSO vehicle control or 20 μM TDCIPP for 1, 2, and 3 days of maturation. Nuclei are marked using DAPI (in blue). Images for oligodendrocytes treated with TMPP and TBPP are shown in Extended Data Fig. 4e. **g,** Quantification of early (O4+), intermediate (O1+), and late (MBP+) oligodendrocytes, after day 1, 2, and 3 of development, normalized to DMSO vehicle control. Data are presented as the mean value ± standard deviation from three biological replicates (OPC batches generated from independent mPSC lines), indicated by closed circle data points. Data for oligodendrocytes treated with TBPP and TMPP are shown in Extended Data Fig. 4f. p-values were calculated using one-way ANOVA with Dunnett post-test correction for multiple comparisons.

To identify at which stage of oligodendrocyte maturation organophosphate flame retardants exert their effect, we cultured developing oligodendrocytes in the presence of TDCIPP, TMPP, and TBPP and assessed maturation over three days. In our toxicity screening platform oligodendrocyte development proceeds through successive stages characterized by expression of known maturation markers (Fig. 3e). Specifically, early oligodendrocytes express the antigen for O4, intermediate oligodendrocytes express the antigen for O1, and mature oligodendrocytes express myelin basic protein (MBP). When we assessed the effects of organophosphate flame retardants on oligodendrocyte generation using immunocytochemistry for early (O4+), intermediate (O1+), and mature (MBP+) oligodendrocytes, we detected a delay in acquisition of O4 expression, and decreased O1 and MBP expression relative to the vehicle (DMSO)-treated negative control at all time points (Fig. 3e-g, and Extended Data Fig. 3e,f). These results suggest that, mechanistically, organophosphate flame retardants arrest the initial progression of early oligodendrocytes to intermediate and mature oligodendrocytes.

### TDCIPP inhibits oligodendrocyte development in human cortical organoids

Next, we used our human cortical organoid model to assess whether organophosphate flame retardants inhibit human oligodendrocyte development. After culturing organoids from day 60 to day 70 in the presence of TDCIPP, we collected samples for immunohistochemistry. In the presence of TDCIPP, an oligodendrocyte marker CC1 was significantly decreased (Fig. 4a,b). Importantly, we found that cell density was unchanged across conditions, suggesting that the deficit in mature oligodendrocytes is indeed a selective inhibition of oligodendrocyte development. Collectively, these results suggest that the presence of TDCIPP is sufficient to arrest human oligodendrocyte development.

**Fig. 4:**
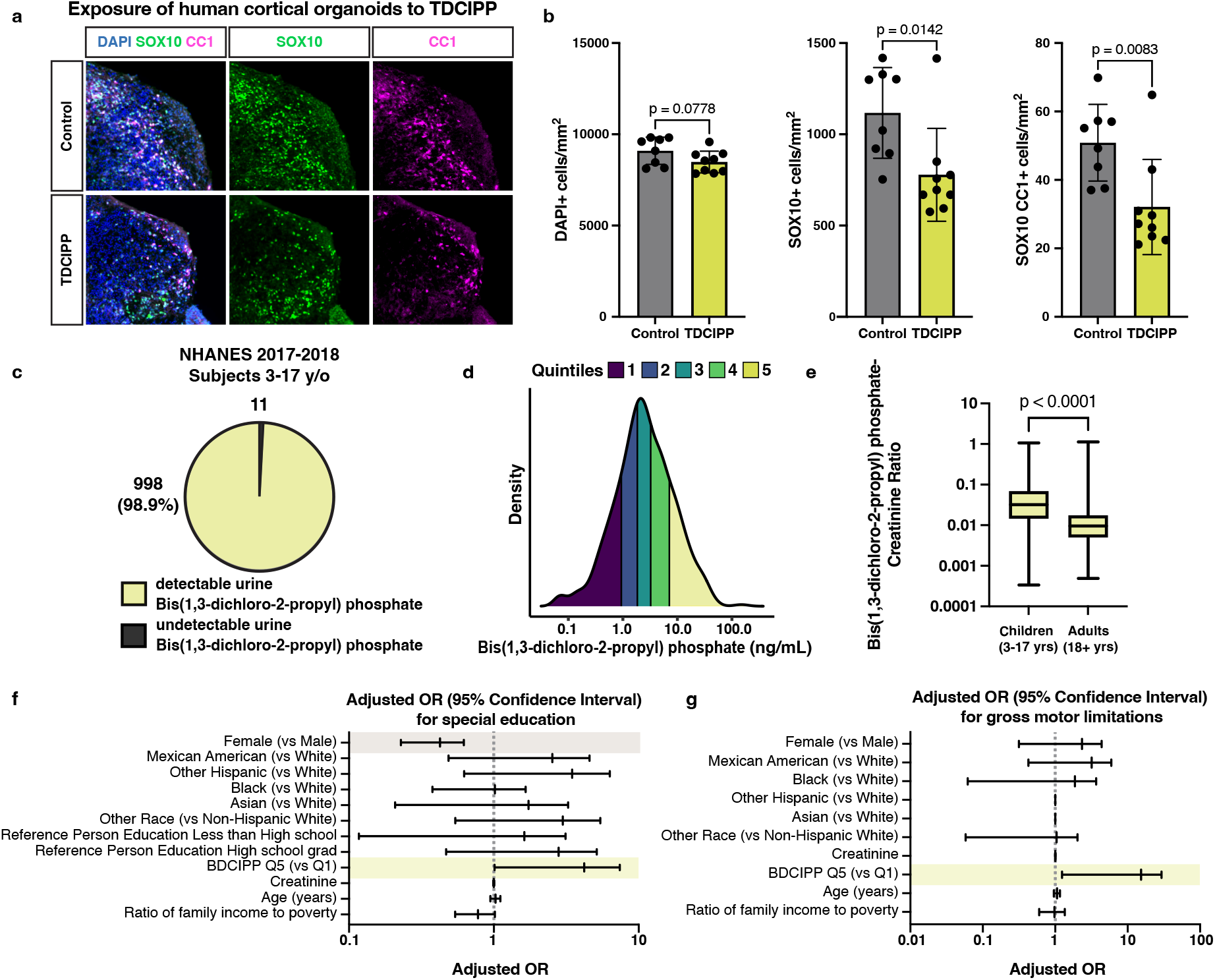
TDCIPP inhibits human oligodendrocyte development and is associated with abnormal neurodevelopmental outcomes in children. **a,** Representative immunohistochemistry images of human cortical organoids treated for 10 days with the flame retardant TDCIPP at 18 μM (approximate IC_75_ in mPSC-derived oligodendrocytes). Images show all cells (DAPI+, in blue), oligodendrocyte lineage cells (SOX10+, in green), and oligodendrocytes (CC1+, in magenta). **b,** Quantification of total cell number (DAPI+ per mm2), oligodendrocyte lineage cells (SOX10+ per mm2) and mature oligodendrocytes (SOX10+CC1+ per mm2) in whole cortical organoids. Data are presented as the mean value ± standard deviation from n ≥ 8 biological replicates (individual organoids) indicated by closed circle data points for TDCIPP-treated organoids. p-values were calculated using Student’s two-tailed t test. **c,** Pie chart showing the number of children ages 3-17 years old from the NHANES 2017-2018 dataset with undetectable and detectable levels of BDCIPP, the urine metabolite of TDCIPP. **d,** Density plot showing the range and quintiles of urine BDCIPP levels in children ages 3-17 years old from the NHANES 2017-2018 dataset. **e,** Boxplot showing creatinine-normalized levels of BDCIPP in children 3-17 years of age and adults aged 18 years and older. p-value was calculated using the Kruskal Wallis oneway ANOVA. **f,** Adjusted odds ratio for the neurodevelopmental outcome: requiring special education or early intervention. Significant odds ratios are highlighted in yellow (BDCIPP Q5 v Q1 OR = 2.7 [95% CI = 1.012-7.407) and gray (Female v Male OR = 0.376 [95% CI = 0.228-0.621]). **g,** Adjusted odds ratio for the neurodevelopmental outcome: gross motor limitations. Significant odds ratios are highlighted in yellow (BDCIPP Q5 v Q1 OR = 6.0 [95% CI = 1.243-29.426).

### TDCIPP exposure during childhood and adolescence is significantly associated with abnormal neurodevelopment

Organophosphate flame retardants are widely used in common products including furniture, building materials, and electronics. Human epidemiological studies investigating potential links between exposure to organophosphate flame retardants and developmental neurotoxicity have largely focused on prenatal exposures^30^. However, given the prolonged period of oligodendrogenesis and myelination after birth, we asked whether postnatal neurodevelopment would be impacted by organophosphate flame retardant exposure throughout childhood and adolescence. To that end, we analyzed data from the US CDC’s National Health and Nutrition Examination Survey (NHANES) to identify levels of childhood exposure to organophosphate flame retardants and associations between exposure and indicators of abnormal cognitive and motor development. The NHANES is a cross-sectional study designed to collect survey, laboratory, and examination data from a nationally representative sample of US children and adults. Due to the complex study design, these data can be leveraged to make estimations, including associations between laboratory-measured exposures and adverse outcomes, that are representative to the entire US population.

Previous work has shown that the urine metabolite bis(1,3-dichloro-2-propy) phosphate (BDCIPP) can accurately estimate exposure to its parent compound, the organophosphate flame retardant TDCIPP^31^. In examining a population of US children 3-17 years of age surveyed between 2017 and 2018 we found that BDCIPP was present in urine samples for 998 out of 1,009, or 98.9%, of children (Fig. 4c,d; Extended Data Fig. 4a). Additionally, the amount of creatinine-adjusted BDCIPP was significantly higher when compared to adults (Fig. 4e). These data indicate that a majority of children were exposed to TDCIPP at the time of the survey and may experience higher internal doses compared to adults.

Using multivariable-adjusted logistic regression, we identified associations between high levels of urinary BDCIPP, and three neurodevelopmental measures: gross motor dysfunction, a need for special education, and a need for mental health services (Extended Data Fig. 4b). These outcomes have been previously used to evaluate neurocognitive and neuromotor function^32,33^. Furthermore, oligodendrogenesis and myelination play essential roles in memory, learning, motor function, and mental health^34–37^. Weighted logistic regression analyses accounted for the NHANES complex survey design and were adjusted for age, sex, race/ethnicity, urine creatinine, as well as socioeconomic confounders previously reported to be associated with each outcome^32,33^. Children in the highest quintile of urinary BDCIPP concentration had increased adjusted odds ratios for all three neurodevelopmental outcomes when compared to children in the lowest quintile of urine BDCIPP concentration. The fully adjusted odds ratios for children in the high BDCIPP concentration group were 2.7 (95% CI = 1.01-7.41) for special education, 4.6 (95% CI = 1.79-12.10) for mental health treatment, and 6.0 (95% CI = 1.24-29.42) for gross motor dysfunction (Fig. 4f,g; Extended Data Fig. 4c). These results indicate that children with high exposure are between 2.7 and 6 times more likely to experience adverse neurodevelopmental outcomes, providing strong evidence of a positive association between organophosphate flame retardant exposure and abnormal neurodevelopment.

## DISCUSSION

Evaluation of chemical safety is essential for the protection of human health. Although the effects of the vast majority of environmental chemicals on the central nervous system are unknown, high throughput screening is a powerful tool to prioritize chemicals of concern based on their toxic effects to physiologically relevant neural cell types and identify environmental triggers of pathogenesis^8,38^. Because oligodendrocytes represent a unique and understudied cell population in developmental neurotoxicology, we developed a toxicity screening platform to interrogate over 1,800 environmental chemicals for their effects on oligodendrocyte development. Through this approach, we identified chemicals from two classes that impede the generation of oligodendrocytes: quaternary compounds and organophosphate flame retardants.

Quaternary compounds are common in personal care products, pharmaceuticals, and anti-static agents. Their prevalent use in disinfectants, including more than half of EPA-registered products for eliminating SARS-CoV-2, is a likely cause of increased human exposure, demonstrated by the doubling in blood levels of some quaternary ammonium compounds since before the COVID-19 pandemic^39,40^. In our oligodendrocyte-specific cytotoxic hits, quaternary compounds as a class were enriched, and in a 3-D model of human prenatal brain development, our data show that these chemicals are specifically cytotoxic to human oligodendrocytes. We report that quaternary compounds induce the integrated stress response in developing oligodendrocytes, resulting in CHOP accumulation and apoptosis. In genetic and inflammatory diseases, developing and regenerating oligodendrocytes are particularly sensitive to ER stress and subsequent prolonged activation of the integrated stress response in part due to the requirement of developing oligodendrocytes to produce large amounts of myelin proteins^41^. This sensitivity may underlie the specific toxicity of quaternary compounds and could initiate or exacerbate pathology in disease.

In developing oligodendrocytes, the IC50 values for quaternary compound toxicity are in the nanomolar range, similar to predicted blood concentrations for many quaternary ammonium compounds in children^42^. Furthermore, we evaluated concentrations as acute exposures; whereas chronic exposures to quaternary compounds spanning oligodendrocyte development could drive toxicity at even lower concentrations, given their capacity for bioaccumulation^39^. When considering *in vitro* cytotoxicity data in mouse and human oligodendrocytes and the potential risk for chronic exposure, the increased use of quaternary ammonium compounds raises significant health concerns for neurodevelopmental toxicity given the ability of quaternary ammonium compounds such as benzalkonium chlorides, found in everyday household disinfecting agents, to pass both the blood placental and the blood brain barriers^43^. We also report the cytotoxicity of one quaternary phosphonium compound, tributyltetradecylphosphonium chloride. Although quaternary phosphonium compounds are less common than their ammonium-based counterparts, they have similar structure and function and may also have increased exposure to humans as they become implemented to combat bacterial resistance seen with quaternary ammonium compounds^25^.

The pervasive use of organophosphate flame retardants has contaminated the environment and increased human exposure, demonstrated by the detection of these chemicals in human blood, urine, breast milk, and cerebrospinal fluid^44,45^. We show that the organophosphate flame retardant TDCIPP arrests the development of mouse oligodendrocytes and inhibits oligodendrocyte generation in human cortical organoids at concentrations similar to estimated blood concentrations in children^46^. Our results and the likelihood of organophosphate flame retardant exposure in children raise potential health concerns as these chemicals may reach higher concentrations in cerebrospinal fluid than blood^47^.

Human epidemiological studies that evaluate prenatal exposure to TDCIPP, have identified associations between maternal exposure and delayed cognitive development^30^. However, these studies focused on solely prenatal exposures. Therefore, the disruption of critical periods of oligodendrogenesis and myelination during neurodevelopment in infancy and childhood by organophosphate flame retardants has yet to be evaluated. We analyzed the US CDC’s NHANES 2017-2018 dataset to identify associations between childhood exposure to organophosphate flame retardants and abnormal neurodevelopmental outcomes. Our logistic regression analyses demonstrate that there are significantly increased odds ratios for children with the highest urinary BDCIPP concentrations for multiple abnormal cognitive and motor outcomes. Continued evaluation of organophosphate flame retardant exposure and direct measurements of white matter development in children would provide critical evidence that chemical-mediated perturbation of oligodendrocyte development influences abnormal cognitive and motor outcomes. Although organophosphate flame retardants are pervasive and human exposure is ubiquitous, behavioral interventions are effective in reducing exposure to TDCIPP and could be considered to minimize potential risks to children^48^.

This work reveals toxicological sensitivities in the oligodendrocyte lineage to common household chemicals and raises potential health concerns for exposure to these chemicals. Continued experimental and epidemiological studies are required to determine the full impact of exposure to quaternary ammonium and phosphonium compounds and organophosphate flame retardants. Results from this study will contribute to the scientific foundation that will inform decisions about regulatory or behavioral interventions designed to reduce chemical exposure and protect human health.

## METHODS

### Induced pluripotent stem cell-derived OPC culture

Mouse OPCs were differentiated from mouse induced pluripotent stem cells (iPSCs) as previously described^21,49^. Briefly, iPSCs were removed from an irradiated mouse embryo fibroblast feeder layer with 1.5 mg/mL collagenase type IV (ThermoFisher, 17104019), dissociated with 0.25% trypsin-EDTA (ThermoFisher, 25200056), and seeded at 7.8×10^4^ cells/cm^2^ on Costar Ultra-Low attachment plates (Sigma, CLS3471). Cells were cultured in media allowing for the expansion and maturation of OPCs for 9 days. On day 10, media was switched to OPC medium, comprised of N2B27 base medium, supplemented with 20 ng/mL FGF-2 (R&D Systems, 233-FB-010), and 20 ng/mL PDGF-AA (R&D Systems, 221-AA). N2B27 base medium consists of Dulbecco’s Modified Eagle Medium/Nutrient Mixture F-12 (DMEM/F-12; ThermoFisher 11320033), supplemented with 1X B-27 Supplement (ThemoFisher, 17504044), 1X N-2 MAX Supplement (ThermoFisher, 17502048), 1X GlutaMAX Supplement (ThermoFisher, 35050079). OPC medium was used over three passages to enrich for OPCs. OPC biological replicates were generated from independent mouse iPSC lines. Mouse iPSC-derived OPCs were used for all experiments unless otherwise noted.

### Primary mouse OPC and astrocyte culture

All animal procedures were performed in accordance with the National Institutes of Health Guidelines for the Care and Use of Laboratory Animals and approved by the Case Western Reserve University Institutional Animal Care and Use Committee. Timed-pregnant mice (C57BL/6N) were ordered from Charles River (Wilmington, MA). Brains from mice were grossly dissected at postnatal day 2 (P2). Cortex tissue was isolated and dissociated using the Miltenyi Tumor Dissociation Kit (Miltenyi, 130-095-929) following the manufacturer’s instructions. Following dissociation, cells were plated in poly-L-ornithine (Sigma, P3655) and laminin (Sigma, L2020) coated flasks and cultured for 24 hours. Culture media consists of N2B27 base medium supplemented with 20 ng/mL FGF-2 (R&D Systems, 233-FB-010), and 50 units/mL-50ug/mL Penicillin-Streptomycin (ThermoFisher, 15070063). After 24 hours of culture, media was switched to astrocyte or OPC enrichment media. OPC enrichment media is comprised of N2B27 base media supplemented with 20 ng/mL PDGF-AA (R&D Systems, 221-AA), 10ng/mL NT-3 (R&D Systems 267-N3), 100 ng/mL IGF (R&D Systems, 291-GF-200), 10 μM cyclic AMP (Sigma, D0260), 100 ng/mL noggin (R&D Systems, 3344NG050) and 50 units/mL-50ug/mL Penicillin-Streptomycin (ThermoFisher, 15070063). OPCs were cultured in this media until the next passage, at which point 50 units/mL-50ug/mL Penicillin-Streptomycin was removed. Astrocyte enrichment media consists of 1:1 DMEM (ThermoFisher, 11960044)—Neurobasal Medium (ThermoFisher, 211-3-49), supplemented with 1X N-2 MAX Supplement (ThermoFisher, 17502048), 1X GlutaMAX Supplement (ThermoFisher, 35050079), 50 units/mL-50ug/mL Penicillin-Streptomycin (ThermoFisher, 15070063), 5 ug/mL N-acetyl cysteine (Sigma, A8199), 10 ng/mL CNTF (R&D 557-NT-010), 5 ng/mL HB-EGF (R&D Systems 259-HE-050), and 20 ng/mL FGF-2 (R&D Systems, 233-FB-010). Media changes were performed every 48 hours and cells were allowed to proliferate, grown to confluency, and either passaged once or cryopreserved. For terminal experiments, astrocytes were thawed and plated into 384-well plates (Perkin Elmer, 6057500) at a density of 4,000 cells per well. Cells were then cultured with maturation media, comprised of 1:1 DMEM and Neurobasal media, supplemented with 1X N-2 MAX, 5 ug/mL N-acetyl cysteine, 1X GlutaMAX Supplement, 1 mM Sodium Pyruvate (ThermoFisher, 11360-070), 5 ng/mL HB-EGF, 10 ng/mL CNTF, 50 ng/mL BMP4 (R&D, 314-BP-050) and 20 ng/mL FGF-2. After 48 hours of culture in astrocyte maturation media, cells were cultured in resting astrocyte media (1:1 DMEM/Neurobasal Medium supplemented with 5 ng/mL HB-EGF) for 72 hours.

### Mouse oligodendrocyte differentiation

OPCs were plated in 96-well plates (Fisher, 167008) coated with poly-L-ornithine (Sigma, P3655) and laminin (Sigma, L2020) at a seeding density of 40,000 cells per well, or 384-well plates coated with poly-D-lysine and laminin (Sigma, L2020) at a seeding density of 12,500 cells per well. Cells were plated in differentiation permissive media, comprised of Dulbecco’s Modified Eagle Medium/Nutrient Mixture F-12 (DMEM/F-12; ThermoFisher 11320033), 1X B-27 Supplement (ThemoFisher, 17504044), 1X N-2 MAX Supplement (ThermoFisher, 17502048), 1X GlutaMAX Supplement (ThermoFisher, 35050079), 10ng/mL NT-3 (R&D Systems 267-N3), 100 ng/mL IGF (R&D Systems, 291-GF-200), 10 μM cyclic AMP (Sigma, D0260), 100 ng/mL noggin (R&D Systems, 3344NG050), and 40 ng/mL T3 (Sigma, T6397) when noted. Cells were differentiated over 3 days and analyzed.

### Immunocytochemistry

Live staining was performed for specific antigens (O1 and O4). Antibodies for O1 and O4 were diluted in N2B27 base medium supplemented with 5% Donkey Serum (v/v) (Jackson ImmunoResearch, 017-000-121) and added to wells for 18 minutes at 37°C. Cells were then fixed with 4% Paraformaldehyde (Electron Microscopy Sciences, HP1-100Kit) for 15 minutes at room temperature, washed with PBS, and incubated overnight at 4°C with primary antibody diluted in PBS supplemented with 5% Donkey Serum (v/v) and 0.1% Triton-X-100 (Sigma, T8787). Primary antibodies included anti-O1 (1:100, CCF Hybridoma core), anti-O4 (1:100, CCF Hybridoma core), and anti-MBP (1:4000, Abcam, ab7349). The following day cells were rinsed with PBS and incubated for 2 hours with the appropriate Alexa Fluor-conjugated secondary antibodies (2 μg/mL, Thermo Fisher) and DAPI (1 μg/mL, Sigma, D8417).

### Chemical screening

Chemicals from the US EPA Toxicity Forecaster (ToxCast) chemical library were obtained through a Material Transfer Agreement with the US EPA. This library contained 1,823 chemicals dissolved in dimethyl sulfoxide (DMSO) at a top target stock concentration of 20 mM (with some chemicals achieving lower stock concentrations based on solubility limits in DMSO) and was stored at −20°C. Screening of the chemical library on OPCs was performed as described previously^21^. CellCarrier Ultra 384-well plates (PerkinElmer, 6057500), pre-coated with poly-D-lysine, were coated with laminin (Sigma, L2020) diluted in N2B27 base media, comprised of Dulbecco’s Modified Eagle Medium/Nutrient Mixture F-12 (DMEM/F-12; ThermoFisher 11320033), 1X B-27 Supplement (ThemoFisher, 17504044), 1X N-2 MAX Supplement (ThermoFisher, 17502048), and 1X GlutaMAX Supplement (ThermoFisher, 35050079). Laminin was dispensed using an EL406 Microplate Washer Dispenser (BioTek) using a 5 μL dispense cassette (BioTek) and incubated for at least 1 hour at 37°C. OPCs were next dispensed in oligodendrocyte differentiation permissive media, at a density of 12,500 cells per well. OPCs were allowed to attach to the plates for 1 hour at 37°C and chemicals were added to plates at a 1:1000 dilution using a Janus automated workstation and 50 nL solid pin tool attachment. Each compound was added at a final test well concentration of 20 μM. Chemicals used for dose response validation were sourced from the primary screening library. DMSO (Sigma, D2650) was added at 1:1000 dilution to negative control wells and 40 ng/mL T3 (Sigma, T6397) was added to positive control wells. After 72 hours, cells were stained with anti-O1 (1:100, CCF Hybridoma core), fixed with 4% Paraformaldehyde (Electron Microscopy Sciences, HP1-100Kit), and imaged using the Operetta High Content Imaging and Analysis system (PerkinElmer).

### Kinetics of oligodendrocyte differentiation

OPCs were seeded at a density of 40,000 cells per well in 96-well plates coated with in poly-L-ornithine (Sigma, P3655) and laminin (Sigma, L2020) in differentiation permissive media supplemented with 40 ng/mL T3 (Sigma, T6397). Cells were treated with TDCIPP (Sigma, 32951), TMPP (Santa Cruz, sc-296611), or TBPP (Millipore, 34188) at a final concentration of 20uM. DMSO (Sigma, D2650), was added at 1:1000 to negative control wells. As described preivously^50^, cells were live stained with anti-O4 (1:100, CCF Hybridoma core), anti-O1 (1:100, CCF Hybridoma core), and fixed with 4% Paraformaldehyde (Electron Microscopy Sciences, HP1-100Kit) after 1-, 2-, and 3-days post-plating. Cells were then stained overnight with anti-MBP (1:4000, Abcam, ab7349) followed by staining with DAPI (1 μg/mL, Sigma, D8417). The Operetta High Content Imaging and Analysis system was used to image 4 fields per well and the percentage of O4, O1, and MBP-positive cells was quantified using the number of DAPI-positive live cells per field.

### High content imaging and quantification

The Operetta High Content Imaging and Analysis system was used to image all 96- and 384-well plates. For each well of the 96- and 384-well plates, 4 fields were captured at 20x magnification. The PerkinElmer Harmony and Columbus software was used to analyze images as described previously^22,50,51^. In brief, nuclei from live cells were identified by DAPI positivity, using thresholding to exclude cell debris or pyknotic nuclei. A region outside of each DAPI-positive nucleus, expanded by 50%, was used to identify oligodendrocytes by positive staining for oligodendrocyte markers (O1 in primary screen and O4, O1, or MBP in kinetics experiments) within this region. Expanded DAPI-positive nuclei that overlapped with O4, O1, or MBP staining were classified as oligodendrocytes. Cell viability was calculated by dividing the number of DAPI-positive nuclei in an experimental well by the average number of DAPI-positive cells in the negative control wells. Oligodendrocyte percentage was calculated by dividing the number of oligodendrocytes by DAPI-positive cells and normalized to negative control wells.

### MTS assay

OPCs were seeded at a density of 12,500 cells per well in CellCarrier Ultra 384-well plates (PerkinElmer, 6057500), pre-coated with poly-D-lysine and laminin (Sigma, L2020) in differentiation permissive media. Cells were allowed to attach for 1 hour at 37°C and compounds were added to plates at a 1:1000 dilution at a final concentration of 20 μM using a Janus automated workstation. DMSO (Sigma, D2650) was added at 1:1000 dilution to negative control wells and 40 ng/mL T3 (Sigma, T6397) was added to positive control wells. Cell viability was assessed after 72 hours using a 3-(4,5-dimethylthiazol-2-yl)-5-(3-carboxymethoxyphenyl)-2-(4-sulfophenyl)-2H-tetrazolium (MTS) assay kit (Abcam, ab197010) according to the manufacturer’s protocol. Absorbance at 490 nm was measured 4 hours after the addition of the MTS dye using a SynergyNEO2 plate reader (BioTek).

### Human cortical organoid production

Human embryonic stem cell research was restricted to *in vitro* culture and *in vitro* cortical organoid generation using the human embryonic stem cell (hESC) line H7 (Wicell, WA07) and was performed following the International Society for Stem Cell Research 2021 Guidelines for Stem Cell Research and Clinical Translation. hESCs were expanded in mTesR1 media (Stem Cell Technologies, 85850) and cortical organoids generated as previously described and with minor modifications^27^. Modifications include replacement of Y-27632 and dorsomorphin with CloneR (Stem Cell Technologies, 5889) and 150nM LDN193189 respectively during the first step in the generation of cortical organoids. For the first 6 days, organoids were cultured with media containing 10 μM SB-43152 (Sigma, S4317) and 150 nM LDN193189, followed by 20 ng/mL EGF (R&D Systems, 236-EG-200) and 20 ng/mL FGF-2 (R&D Systems, 233-FB-010) on days 7 to 25. On days 27 to 40 organoids were fed on alternate days with media containing 20 ng/mL NT-3 (R&D Systems, 267-N3) and 10 ng/mL BDNF (R&D Systems 248-BD). To expand OPC populations, 10 ng/mL PDGF-AA (R&D Systems, 221-AA) and 10 ng/mL IGF (R&D Systems, 291-GF-200) were added to organoid cultures every other day between days 51 and 60. To induce the differentiation of oligodendrocytes 40 ng/mL T3 (Sigma, T6397) was added on alternate days between days 60 to 70. Organoids were treated with DMSO or chemicals beginning on day 60 and harvested on day 70. Methyltrioctylammonium chloride and tributyltetradecylphosphonium chloride were sourced from the primary screening library and added at their IC_90_ concentrations. TDCIPP (Sigma, 32951) was added at its approximate IC_75_ concentration.

### Cortical organoid immunohistochemistry

Cortical organoids were treated on alternating days between days 61 to 70 with quaternary ammonium and phosphonium compounds or organophosphate flame retardants at their approximate IC_90_ and IC_75_ concentrations respectively. Organoids were harvested on day 70, washed in PBS, and fixed overnight with ice-cold 4% Paraformaldehyde (Electron Microscopy Sciences, HP1-100Kit). On the following day organoids were washed with PBS and cryoprotected using a 30% sucrose solution. Organoids were then embedded in OCT and sectioned at 15 μM. Slides were washed with PBS and incubated overnight with anti-SOX10 (1:200, R&D, AF2864) and anti-APC CC1 (1:200, Millipore, MABC200), followed by labeling with Alexa Fluor-conjugated secondary antibodies (2 μg/mL, Thermo Fisher). Slides were imaged at 10x magnification using a Hamamatsu Nanozoomer S60. Quantification of positive cells was performed using QuPath software (https://qupath.github.io/)^52^.

### Cell death inhibitor testing

OPCs were seeded in 384-well plates (PerkinElmer, 6057500) pre-coated with poly-D-lysine and laminin (Sigma, L2020) at a density of 12,500 cell per well and allowed to attach for 1 hour at 37°C. Cell death inhibitors quinoline-Val-Asp-Difluorophenoxymethylketone (QVD-OPH) (Selleck, S7311), ferrostatin-1 (Selleck, S7243), and necrostatin-1 (Selleck, S8037), were added using a Janus automated workstation and 50 nL solid pin tool attachment in 8-point dose response (80 nM to 10 μM), and incubated for 1 hour at 37°C. Methyltrioctylammonium chloride or tributyltetradecylphosphonium chloride was added to all wells at IC_90_ concentrations (approximately 100 nM), and oligodendrocytes were allowed to develop for 72 hours. Negative control wells contained only methyltrioctylammonium chloride or tributyltetradecylphosphonium chloride. Positive control wells contained vehicle (DMSO). Cells were fixed with 4% Paraformaldehyde (Electron Microscopy Sciences, HP1-100Kit) and stained with and DAPI (1 μg/mL, Sigma, D8417). Imaging was performed with the Operetta High Content Imaging and Analysis system (PerkinElmer) and the PerkinElmer Harmony and Columbus software was used to quantify DAPI-positive nuclei.

### RNA sequencing

OPCs were plated in 6-well plates coated with poly-L-ornithine (Sigma, P3655) and laminin (Sigma, L2020) in OPC medium (Fisher Scientific, 14-832-11) and allowed to attach for one hour. OPCs were incubated with methyltrioctylammonium chloride and tributyltetradecylphosphonium chloride at their approximate IC_90_ concentrations for 4 hours. OPCs were then lysed in TRIzol (Invitrogen, 15596018) and RNA was extracted by phenol-chloroform extraction and purified using the RNeasy Mini Kit (Qiagen, 74104). Samples were sent to Novogene for library preparation and mRNA sequencing. Libraries were generated according to protocols from the NEBNext Poly(A) mRNA Magnetic Isolation Module (NEB, E7490L) and NEBNext Ultra RNA Library Prep Kit for Illumina (NEB, E7530L) and then evenly pooled and sequenced on the Illumina NovaSeq with 150bp paired-end reads and a read dept of at least 20 million reads per sample. Salmon 1.8.0. (https://github.com/COMBINE-lab/salmon)^53^ was used to align reads to the mm10 genome and quantify transcript abundance as transcripts per million (TPM) values. The R package tximport was used to convert TPM values into gene-TPM abundance matrices.

### qRT-PCR

OPCs were seeded at a density of 1,000,000 cells per well in poly-L-ornithine (Sigma, P3655) and laminin (Sigma, L2020) 6-well plates. OPCs were lysed using TRIzol (Invitrogen, 15596018) and RNA was isolated as described for RNA sequencing. RNA quantity and quality was assessed using a NanoDrop spectrophotometer and cDNA was synthesized using an iScript cDNA Synthesis Kit (Biorad, 1708891) following the manufacturer’s instructions. qRT-PCR was performed using TaqMan gene expression assays (Thermo Fisher, 4369016) and run on an Applied Biosystems QuantStudio 3 real-time PCR system. *Rpl13a* (Mm05910660_g1) was used as an endogenous control and probes for *Ddit3* (Mm01135937_g1) were normalized to the endogenous control.

### Gene set enrichment analysis

Gene set enrichment analysis (GSEA) software was used to calculated normalized enrichment scores, in hallmark datasets using 1000 gene-set permutations, classical scoring, and signal-to-noise metrics (https://www.gsea-msigdb.org/gsea/index.jsp). GSEA software generated normalized enrichment scores and false discovery rates. The integrated stress response gene set was curated from two published gene sets (Supplementary Table 3)^54,55^.

### US EPA ToxCast data

Data from the US EPA ToxCast invitroDBv3.3, was used to assign use categories to chemical hits and obtain median cytotoxicity values. Chemical use categories were assigned based on “collected data on functional use” or “products use categories” obtained from the CompTox dashboard (https://comptox.epa.gov/dashboard/). Cytotoxicity median values were generated using data from the invitroDBv3.3 and the R package tcpl (ToxCast Analysis Pipeline)^56^.

### ToxPrint chemotype enrichment analysis

ToxPrint chemotype enrichment analysis to identify enriched ToxPrints was performed as previously described using the publicly available ToxPrint feature set (https://toxprint.org/) and Chemotyper visualization application (https://chemotyper.org/)^56^. In separate analyses, chemical sets of interest were assigned as the “positive” chemical set and the remaining chemicals were assigned as the “negative” chemical set. Fischer’s exact test was used to calculate p values for enriched ToxPrints in the positive chemical set compared to the negative set. Odds ratios were calculated as described previously^57^. ToxPrints had a p value < 0.05 and odds ratio > 3 were considered significant and analyzed further.

### Statistical analysis

GraphPad prism was used to perform curve-fitting for the calculation of IC50, IC_75_ and IC_90_ values and all statistical analyses unless otherwise specified. Data are typically presented as the mean ± standard deviation (SD) as described in figure legends. A p value < 0.05 was considered statistically significant. For box-and-whisker plot, the box extends from 25^th^ to 75^th^ percentiles, with a line in the middle of the box at the median. Whiskers extend from the minimum to maximum values in the dataset.

### NHANES data source and study population

Anonymized data from the National Health and Nutrition Examination Survey (NHANES) were downloaded from the US Centers for Disease Control and Prevention (CDC) website (https://www.n.cdc.gov/nchs/nhanes). The 2017-2018 NHANES was approved by the CDC’s National Center for Health and Statistics Ethics Review Board (protocol #2018-01) and is provided as anonymized data for public download. NHANES data were downloaded from the 2017-2018 dataset and used for logistic regression analyses.

### NHANES exposure assessment

Exposure to tris(1,3-dichloro-2-propyl) phosphate (TDCIPP) was assessed based on urinary concentrations of bis(1,3-dichloro-2-propyl) phosphate (BDCIPP). Levels of urine BDCIPP were measured in the 2017-2018 dataset in all children ages 3-5 years of age and a one-third subset of children ages 6-17 years of age. Detailed methods for the measurement of BDCIPP urine concentration are provided online via the NHANES website (https://www.n.cdc.gov/nchs/nhanes). Briefly, BDCIPP concentration was determined by solid phase liquid extraction followed by isotope dilution-ultrahigh performance liquid chromatography-tandem mass spectrometry. The lower limit of detection (LLOD, in ng/mL) for this assay was 0.1. For analytes below the LLOD, an imputed fill value was generated by dividing the LLOD by the square root of 2.

### NHANES outcomes

Three neurodevelopmental outcomes, including reported special education, gross motor impairment, and mental health treatment, were assessed. Children greater than or equal to 16 years of age were interviewed directly. A proxy provided answers to questions for children below age 16 years of age. Proxies were asked the following questions to assess motor dysfunction: “Does Sample Person (SP) have an impairment or health problem that limits his/her ability to walk/run/play?”. Children over the age of 16 were asked “Do you have an impairment or health problem that limits your ability to walk/run?” For assessment of special education utilization children and proxies were asked “Does SP receive Special Education or Early Intervention Services?”. Determining whether a child required mental health services, children and proxies were asked “During the past 12 months, have you/has SP seen or talked to a mental health professional such as a psychologist, psychiatrist, psychiatric nurse, or clinical social worker about your/his/her health?”

### NHANES covariates

NHANES collects data on other covariates including demographic and socioeconomic information. Subject age was determined based on participant’s date of birth. Participant’s gender was queried at the time of the survey. Race/ethnicity was divided into five categories: Mexican American, other Hispanic, non-Hispanic white, non-Hispanic black, non-Hispanic Asian, other race-including multi-racial. Education level of the household reference person was split into three categories: less than high school degree, high school grad/GED or some college/AA degree, and college graduate or above. Poverty income ratios were calculated by dividing family income to poverty guidelines determined by the Department of Health and Human Services for the given survey year. Ratios at or above 5.00 were coded as 5.0. Department of Health and Human Services poverty guidelines were utilized to determine the ratio of family income to poverty. Detailed information on all NHANES covariates is available online (https://www.n.cdc.gov/nchs/nhanes).

### NHANES Statistical analysis

All analyses were performed using SPSS. The complex samples logistic regression procedure in SPSS was used to perform multivariable-adjusted logistic regression to estimate adjusted odds ratios and 95% confidence intervals. All analyses specified strata, cluster, and environmental weight variables to account for the NHANES complex survey design. To assess the presence of non-linear relationships between urinary BDCIPP and neurodevelopmental outcomes, BDCIPP was evaluated in quintiles. We constructed two main models containing the following covariates: urinary creatinine, sex, age, race/ethnicity, ratio of family income to poverty, and education level of the household reference person. Age, urinary creatinine, and ratio of family income to poverty were modeled as continuous variables while sex, race/ethnicity, and education level of the household reference person were included as discrete covariates.

## DATA AVAILABILITY

Primary screening results are available in Supplementary Table 1 and will be included in the next public release of the US EPA ToxCast database. RNA-seq datasets generated in this study have been deposited in Gene Expression Omnibus (https://www.ncbi.nlm.nih.gov/geo/) under accession code GSE212190. Access key: qhwrmiisxdqxvgd

## ACKNOWLEDGEMENTS

This work was supported by grants from the National Institutes of Health R35NS116842 (P.J.T.), F31NS124282 (E.F.C.), T32NS077888 (E.F.C.), and T32GM007250 (E.F.C.). B.L.L.C. is supported by a NMSS Career Transition Fellowship. Institutional support was provided by CWRU School of Medicine and philanthropic support was generously contributed by the Fakhouri, Long, Walter, Peterson, Goodman, and Geller families. Additional support was provided by the Small Molecule Drug Development and Light Microscopy Imaging core facilities of the CWRU Comprehensive Cancer Center (P30CA043703). The US Environmental Protection Agency provided the ToxCast screening library through MTA with CWRU and supported the effort of EPA employees (T.J.S. and K.P-F.). We are grateful to D. Adams, A. Wynshaw-Boris, K. Carr, K. Lee, J. Kristell, M. Scavuzzo, K. Allan, and A. Gartley for technical assistance and/or discussion.

## DISCLAIMER

This work was supported in part by the US Environmental Protection Agency and has been reviewed and approved for publication by the US EPA’s Center for Computational Toxicology and Exposure. Approval for publication does not signify that the contents reflect the views of the Agency, nor does mention of trade names or commercial products constitute an endorsement or recommendation for use.

## AUTHOR CONTRIBUTIONS

E.F.C., B.L.L.C., T.J.S., and P.J.T. conceived this study to screen effects of environmental chemicals on oligodendrocyte development. E.F.C., B.L.L.C., and P.J.T. designed and managed the experimental studies. E.F.C, B.L.L.C, and S.Y. performed, quantified, and analyzed in vitro experiments using mouse OPCs including primary screening, and immunocytochemistry. E.F.C. and S.Y. performed dose-curve validations and qPCR. B.L.L.C. isolated mouse astrocytes and performed primary screening for astrocytes. E.F.C. performed RNA-seq analysis. K.P-F performed ToxPrint chemotype enrichment analyses and T.J.S. and K.P.F. guided categorization of chemical screen hits. E.F.C. designed and performed linear regression analyses using data from the National Health and Nutrition Examination Survey. M.M. and E.F.C. performed cortical organoid experiments. Y.F. managed the chemical library and pipelined primary screening data. E.F.C assembled all figures. E.F.C. and P.J.T. wrote the manuscript with input from all authors.

---

**Extended Data Fig. 1:**
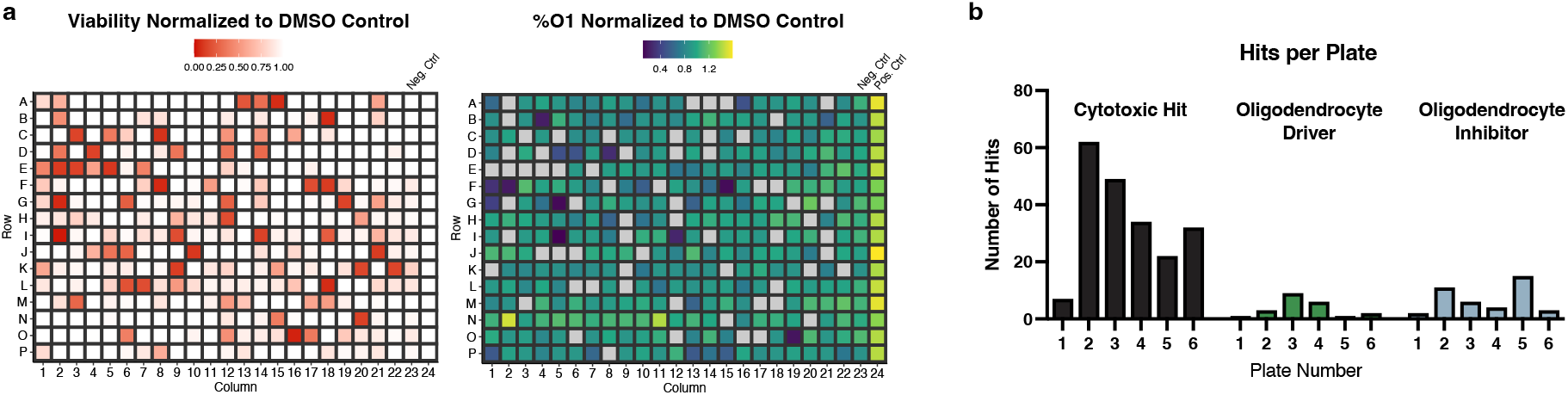
Screening a library of environmental chemicals in developing oligodendrocytes identifies cytotoxic chemicals and modulators of oligodendrocyte generation. **a,** Representative heatmaps of one of six primary screening 384-well plates depicting cytotoxic compounds (red), oligodendrocyte inhibitors (blue), and drivers (green). Viability and percent O1+ oligodendrocytes are normalized to vehicle control (DMSO). Thyroid hormone, a known driver of oligodendrocyte generation, is included as a positive control for oligodendrocyte development. **b,** Quantification of hits across 6 primary screening plates showing distribution of chemicals identified as cytotoxic (black), drivers (green), and inhibitors (blue).

**Extended Data Fig. 2:**
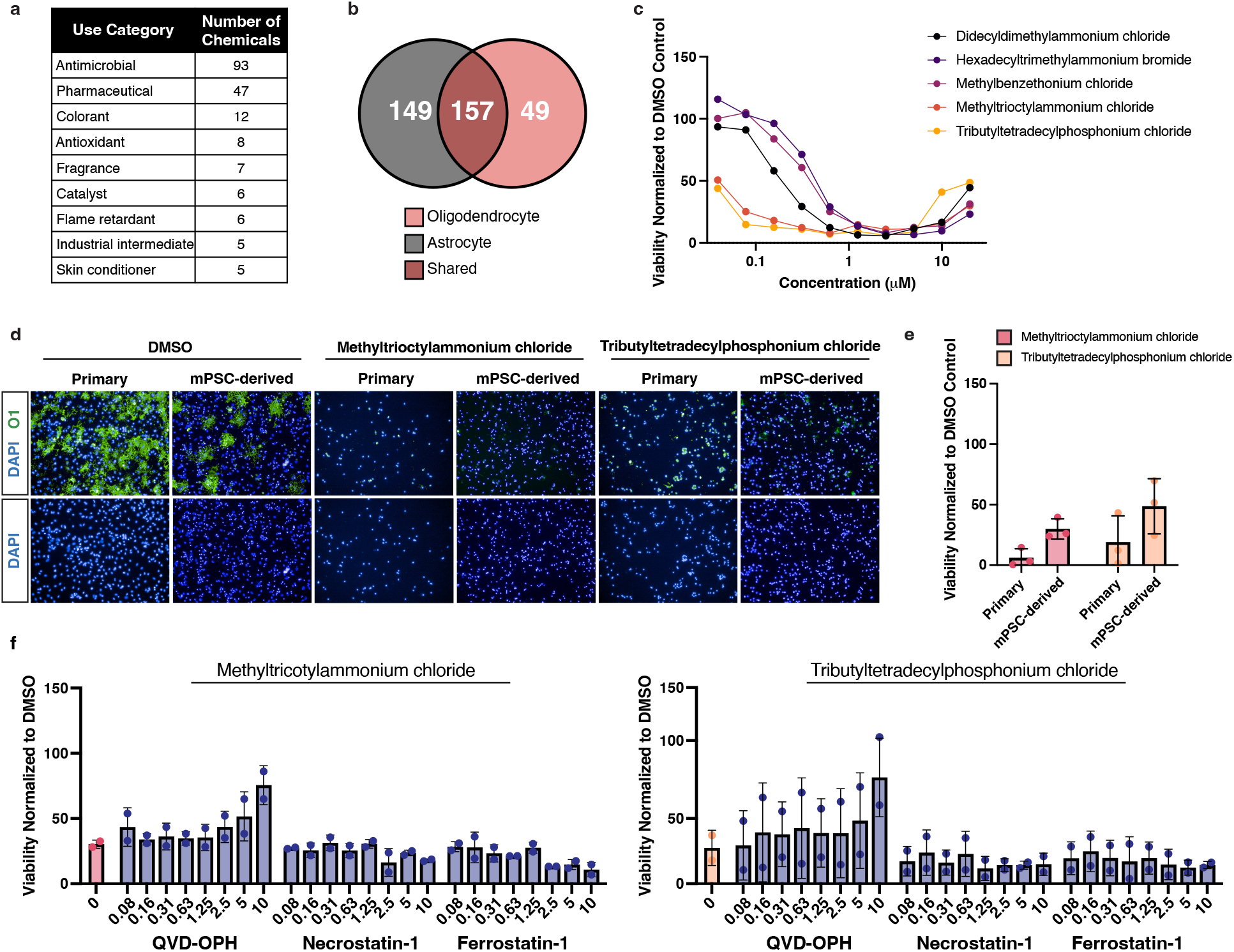
Quaternary compounds are specifically cytotoxic to oligodendrocyte development and induce apoptosis. **a,** Table of the top use categories for the 206 validated cytotoxic chemicals and the number of chemicals belonging to each category. **b,** Venn diagram showing the overlap of 206 validated cytotoxic chemicals identified in the oligodendrocyte screen (in red) compared to cytotoxic hits identified in an identical screen performed in mouse astrocytes (in gray). **c,** Quaternary compounds tested in 10-point dose response from (40 nM to 20 μM), on developing oligodendrocytes quantifying cell number (DAPI+). Data are presented as the mean value from 3 biological replicates (OPC batches generated from independent mPSC lines). **d,** Representative immunohistochemistry images of mPSC-derived oligodendrocytes (O1+, in green) and primary mouse oligodendrocytes treated with methyltrioctylammonium chloride and tributyltetradecylphosphonium chloride. Nuclei are marked by DAPI (in blue). **e,** Quantification of viability of mouse primary and PSC-derived oligodendrocytes treated with 20μM methyltrioctylammonium chloride or tributyltetradecylphosphonium chloride. Data are presented as the mean ± standard deviation from three biological replicates, represented by closed circles (from independent primary OPC isolations or OPC batches from independent mPSC lines). **f,** Quantification of oligodendrocyte viability normalized to DMSO control. Developing oligodendrocytes were cultured for 3 days in the presence of 120 nM methyltrioctylammonium chloride or 100 nM tributyltetradecylphosphonium chloride at (approximate IC_75_ in mPSC-derived OPCs), and cell death inhibitors QVD-OPH, necrostatin-1, and ferrostatin-1, in 8-point dose response (80 nM to 10 μM). Data are presented as the mean ± standard deviation from two biological replicates, represented by closed circles (OPC batches generated from independent mPSC lines).

**Extended Data Fig. 3:**
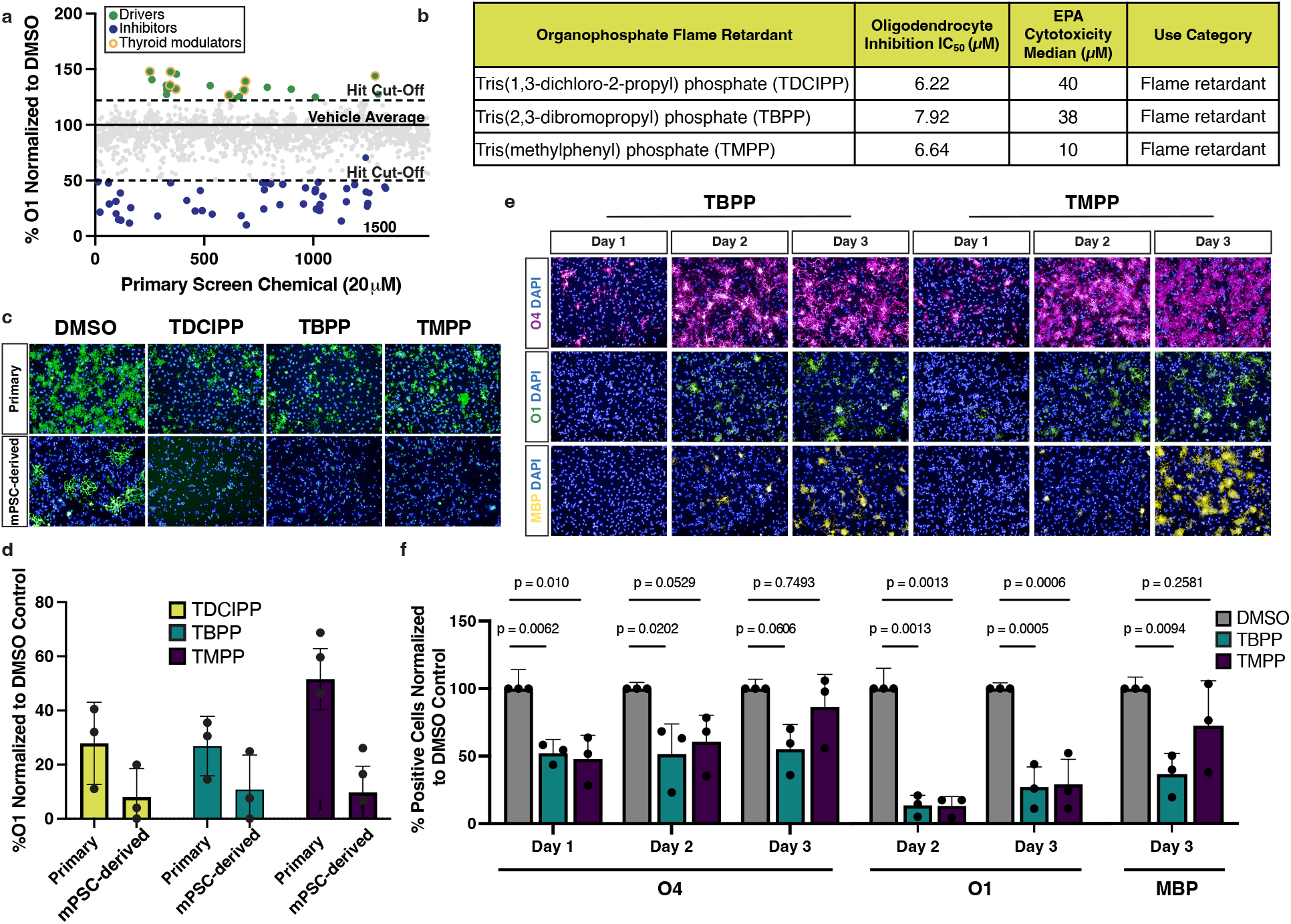
Organophosphate flame retardants inhibit oligodendrocyte development. **a,** Primary chemical screen of 1,539 non-cytotoxic environmental chemicals showing the effect of individual chemicals on oligodendrocyte generation, presented as percent O1 + cells normalized to the DMSO control, as shown in Fig. 2a. Two dotted lines show the hit cutoffs for identification of oligodendrocyte drivers and inhibitors. Drivers result in an increase of O1+ percentage by 22% (>3 standard deviations) compared to negative DMSO control. Inhibitors reduce O1+ percentage by more than 50% (>7 standard deviations) compared to negative DMSO control. Thyroid modulators are highlighted in yellow. **b,** Table shows IC50 concentrations, cytotoxicity median values, and use categories for three organophosphate esters identified as inhibitors of oligodendrocyte development. **c,** Representative immunohistochemistry images of oligodendrocytes, generated from mPSC-derived OPCs and mouse primary OPCs, tested with three organophosphate flame retardants at 20 μM. Generation of oligodendrocytes was evaluated using the oligodendrocyte marker O1 (green). Nuclei are marked with DAPI (in blue). **d,** Quantification O1+ mPSC-derived and primary oligodendrocytes, shown as a percentage of DAPI+ cell number, across three biological replicates, represented at closed circles (independent isolations of primary OPCs and OPC batches generated from independent mPSC lines). **e,** Immunohistochemistry images of early (O4+, in magenta), intermediate (O1+, in green), and late (MBP+, in yellow) oligodendrocytes treated with 20 μM TBPP or TMPP. Control images and TDCIPP treated oligodendrocytes are shown in Fig. 2e. Nuclei are marked with DAPI (in blue). **f,** Quantification of primary oligodendrocytes at the early (O4+), intermediate (O1+), and late (MBP+) stage, shown as a percentage of DAPI+ cell number, over three days of development. Data are presented as the mean ± standard deviation from three biological replicates (OPC batches generated from independent mPSC lines), indicated by closed circle data points. p-values were calculated using one-way ANOVA with Dunnett post-test correction for multiple comparisons.

**Extended Data Fig. 4:**
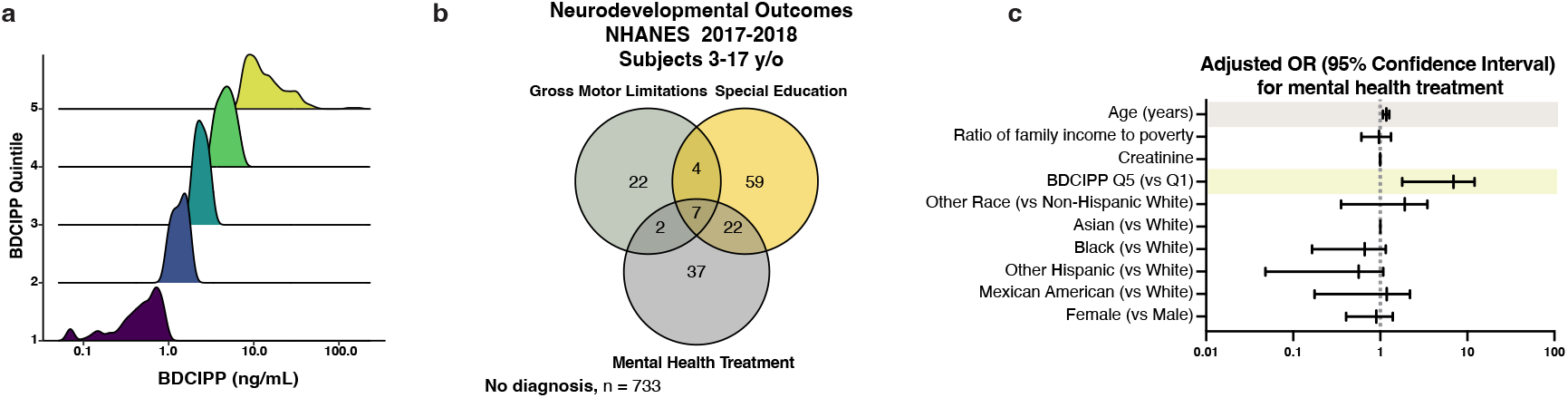
TDCIPP is associated with abnormal neurodevelopmental outcomes in children. **a,** Venn diagram showing co-occurrence of three neurodevelopmental outcomes in the study population. **b,** Density plots showing the distribution of urine BDCIPP levels within individual quintiles. **c,** Adjusted odds ratio for the neurodevelopmental outcome “sought mental health treatment”. Significant odds ratios are highlighted in blue (BDCIPP Q5 v Q1 OR = 4.6 [95% CI = 1.785-12.104) and significant covariates are highlighted in gray (p < 0.001).

## Supplementary Information Tables (provided as separate .xlsx files)

**Supplementary Table 1. Primary screening results**

Primary screening results showing the effects of environmental chemicals on the viability and generation of developing oligodendrocytes. Cytotoxic chemicals, identified by comparing DAPI-positive cell number in treated wells to vehicle (DMSO), are included in an additional sheet. Non-cytotoxic chemicals were assessed for effects on oligodendrocyte development. Chemicals that increased or decreased oligodendrocyte number, measured by the percentage of O1-positive oligodendrocytes, were identified as drivers or inhibitors, and are included in separate sheets.

**Supplementary Table 2. ToxPrint enrichment analysis**

Computational analysis showing cytotoxicity-associated structures assigned to all chemicals in the 1,823 chemical library, and additional sheets for enriched structures within the top cytotoxic hits and inhibitors or oligodendrocyte development.

**Supplementary Table 3. Expression data for quaternary compound treated OPCs**

Gene set enrichment results for OPCs treated with methyltrioctylammonium chloride or tributyltetradecylphosphonium chloride for 4 hours at IC_90_ concentrations.

